# Gene Set Overlap: An Impediment to Achieving High Specificity in Over-representation Analysis

**DOI:** 10.1101/319145

**Authors:** Farhad Maleki, Anthony J. Kusalik

**Author notes:** This paper was published in Bioinformatics 2019. Please cite it as follows: Maleki, Farhad, and Anthony J. Kusalik. “Gene set overlap: An impediment to achieving high specificity in over-representation analysis.” Proceedings of the 12th International Joint Conference on Biomedical Engineering Systems and Technologies. Vol. 3. 2019.

## Abstract

Gene set analysis methods are widely used to analyze data from high-throughput “omics” technologies. One drawback of these methods is their low specificity or high false positive rate. Over-representation analysis is one of the most commonly used gene set analysis methods. In this paper, we propose a systematic approach to investigate the hypothesis that gene set overlap is an underlying cause of low specificity in over-representation analysis. We quantify gene set overlap and show that it is a ubiquitous phenomenon across gene set databases. Statistical analysis indicates a strong negative correlation between gene set overlap and the specificity of over-representation analysis. We conclude that gene set overlap is an underlying cause of the low specificity. This result highlights the importance of considering gene set overlap in gene set analysis and explains the lack of specificity of methods that ignore gene set overlap. This research also establishes the direction for developing new gene set analysis methods.

## 1 INTRODUCTION

High-throughput “omics” technologies have been widely used to investigate biological questions that require screening of a large number of biomolecules. The main challenge facing these technologies is analyzing the generated data to gain biological insight. An RNA-Seq experiment, for example, may suggest several hundred genes as being differentially expressed. Manual interpretation of such a large set of genes is impractical and susceptible to investigator bias toward a hypothesis of interest.

Gene set analysis is a well-established computational approach to gain biological insight from data resulting from high-throughput gene expression experiments (Huang et al., 2009). It relies on the assumption that most biological processes are the consequence of a coordinated activity of a group of genes. Therefore, the primary goal of gene set analysis is to detect concordant changes in expression patterns of predefined groups of genes, referred to as gene sets. Members of a given gene set often share a common biological function or attribute. MSigDB (Liberzon et al., 2011), GeneSigDB (Culhane et al., 2011), GeneSetDB (Araki et al., 2012), Go-Elite (Zambon et al., 2012), and Enrichr (Kuleshov et al., 2016) are among the most widely used gene set databases. These databases have been generated from various sources including GO (Ashburner et al., 2000), KEGG (Kanehisa et al., 2016), Reactome (Joshi-Tope et al., 2005), and BioCarta (Nishimura, 2001).

Often gene set analysis methods report a large number of gene sets as being differentially enriched, where the majority of the reported gene sets are biologically irrelevant or uninformative (Tarca et al., 2013). The rapid growth of the size of gene set databases is intensifying this issue. Consequently, gaining biological insight from the results of gene set analysis is becoming more challenging and prone to investigator biases in favour of a hypothesis of interest. For example, Araki et al. used GeneSetDB to analyze a list of 79 differentially expressed Affymetrix probe sets (Araki et al., 2012) resulting from an experiment where endothelial cells were induced to undergo apoptosis (Johnson et al., 2004). After correction for multiple hypothesis testing, they reported 1694 gene sets as statistically significant, i.e. differentially enriched. Interpreting this large number of gene sets is challenging.

Understanding the factors contributing to low specificity in gene set analysis helps in choosing methods that are more robust against these factors. Such an understanding also facilitates interpreting the results of gene set analysis methods and accelerates the development of new methods that address these contributing factors to achieve higher specificity without sacrificing sensitivity and accuracy.

Specificity of gene set analysis methods in the absence of differential expression of genes has been studied. Tarca et al. (Tarca et al., 2013) investigated the specificity of sixteen gene set analysis methods in the absence of differential expression and showed that even when there is no differential expression, some gene set analysis methods produce a large number of false positives. However, their approach cannot be used to assess the specificity of a gene set analysis method in the presence of differentially expressed genes.

Overlap between gene sets has been suggested as being responsible for the low specificity of gene set analysis methods. To deal with overlap between gene sets, PADOG (Tarca et al., 2012) assigns lower weights to genes that belong to more than one gene set. For a given gene *g*, this weight is negatively correlated with the number of gene sets containing *g*. TopGO (Alexa et al., 2006) is another attempt to deal with gene set overlap. It considers that Gene Ontology (GO) terms are organized as a directed acyclic graph encoding a hierarchy of general-to-more-specific terms. This structure leads to commonality between the genes corresponding to a child node and those of its parent(s). TopGO proposes a gene elimination and a gene down-weighting procedure to decorrelate the GO graph structure resulting from these relations. MGSA (Bauer et al., 2010) utilises a Bayesian approach that considers the overlap between GO categories to reduce the number of false positives. SetRank (Simillion et al., 2017) is another attempt at reducing the number of false positives by considering the overlap between gene sets.

Parallel to the development of gene set analysis methods, various gene set databases have been developed. The prevailing trend in developing gene set databases has been introducing more gene sets and increasing database size. Figure 1 illustrates the growth of MSigDB across its versions. This gene set database has been designed for gene set analysis in human, and its current version includes gene sets from various sources such as GO, KEGG, Reactome, and BioCarta. This gene set database has undergone a 13-fold increase in the number of gene sets compared to its first version. Given the limited number of known genes for human, this steep growth leads to an increase in the number of gene sets overlapping with each other.

**Figure 1:**
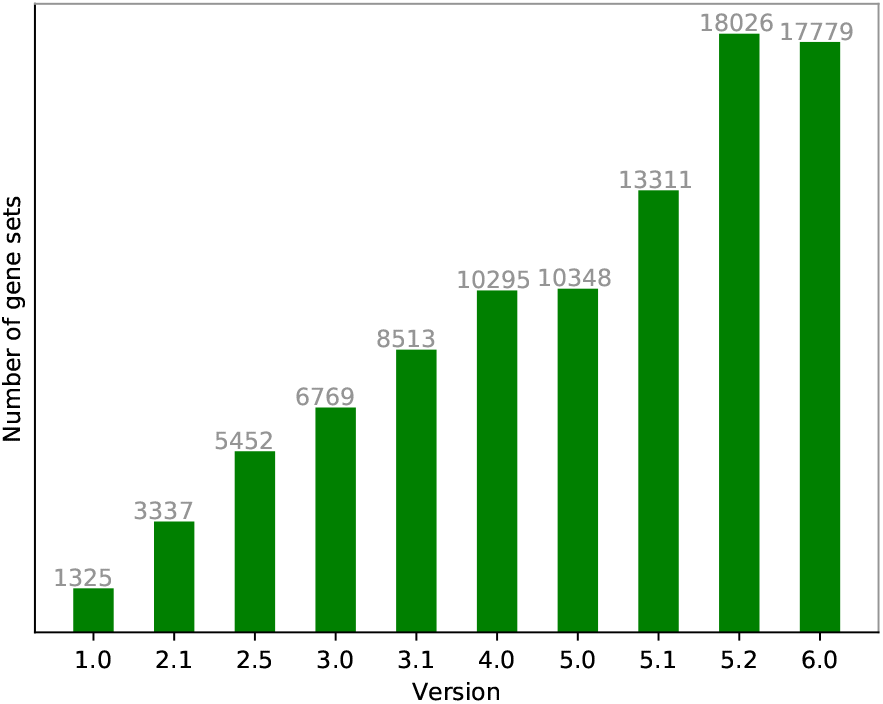
The number of gene sets in different versions of MSigDB

To the best of our knowledge, there is no systematic study of the effect of gene set overlap on the results of gene set analysis. In this paper, we investigate the hypothesis that gene set overlap plays a prominent role in the lack of specificity of over-representation analysis (ORA), which is one of the most widely used gene set analysis methods (Draghici et al., 2003).

The rest of the paper is organised as follows. In Section 2, we briefly describe ORA. In Section 3, we show that gene set overlap is a ubiquitous phenomenon in gene set databases; we use quantitative measures to visualize gene set overlap in GeneSetDB (Araki et al., 2012), GeneSigDB (Culhane et al., 2010), and MSigDB (Liberzon et al., 2011), which are well-established gene set databases. In Section 4, using these quantitative measures, we introduce a methodology to study the effect of gene set overlap on the specificity of ORA. In Section 5, we describe the experimental results; using the methodology introduced in Section 4, we statistically investigate the effect of gene set overlap on the specificity of ORA by assessing the correlation between gene set overlap and specificity. In Section 6, we discuss the implication of gene set overlap and the challenges it entails. We also provide suggestions for developing and evaluating gene set analysis methods. Finally, Section 7 offers a short summary and conclusion.

## 2 OVER-REPRESENTATION ANALYSIS

Many algorithms have been proposed and used for gene set analysis, of which ORA is one of the most widely used. Due to its simplicity, well-established underlying statistical model, and ease of implementation, ORA is available through many tools (Beißbarth and Speed, 2004), (Berriz et al., 2003), (Boyle et al., 2004), (Jiao et al., 2012), (Maere et al., 2005), (Wang et al., 2017), (Wrobel et al., 2005), (Young et al., 2005), (Zeeberg et al., 2003), (Zeeberg et al., 2005), (Zhang et al., 2005). This method defines a concordant change in expression pattern of members of a given gene set as a change that is unlikely to happen by chance. It also quantifies the concept of change as the number of differentially expressed genes in a pairwise comparison of phenotypes, e.g. “cancerous” versus “non-cancerous”.

ORA can be outlined as follows (Drăghici et al., 2003). Suppose that data analysis for an experiment using a high-throughput technology predicts a set of differentially expressed genes *L*, and that the intersection of *L* and a given gene set *G_i_* contains 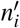 genes. In addition, assume that the set of background genes, i.e. all genes with a non-zero probability of being differentially expressed, contains *n* genes. For example, the background genes in a microarray study can be the set of all genes represented on the arrays. Denote the background set as *U*. Let 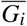 refer to the complement of *G_i_* with respect to *U*, i.e. all genes in *U* but not in *G_i_*. Given *L*, *G_i_*, and *U*, ORA assesses whether the number of differentially expressed genes in *G_i_* is more than what it should be just by chance, i.e. it is over-represented. Table 1 represents ORA as a contingency table, where ‖ • ‖ is the cardinality operator.

**Table 1:**
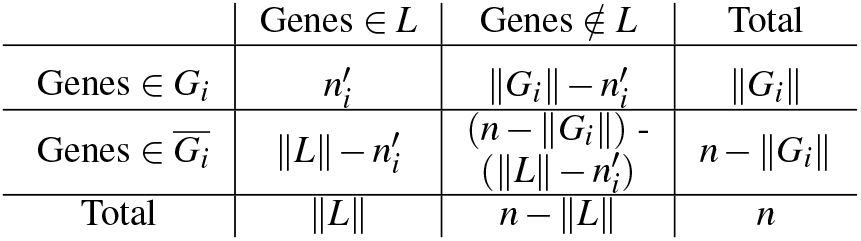
Representation of ORA as a contingency table. Each cell contains a count of genes satisfying the condition given by the row and column.

Assuming that genes are selected using a simple random sampling approach, ORA can be modeled using a hypergeometric distribution (Drăghici et al., 2003). Accordingly, the probability of having 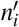 genes from *G_i_* among differentially expressed genes, i.e. *L*, is as follows:

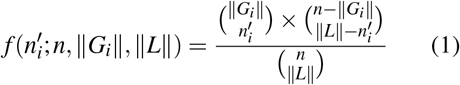

In addition, Fisher’s exact test can be used to examine the significance of the association between genes in *G_i_* and genes in *L*. The *p*-value can be calculated for over-representation of *G_i_* based on Equation 2.

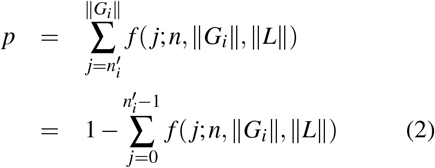

## 3 Overlap in gene set database

ORA, as with other gene set analysis methods, relies on availability of a gene set database. Gene set databases are developed by collecting genes that are manually or computationally inferred to share a common biological function or attribute. The availability of *a priori* knowledge through public repositories such as GO (Ashburner et al., 2000), KEGG (Kanehisa et al., 2016), and OMIM (Hamosh et al., 2002) makes it possible to develop gene set databases. There are many publicaly available gene set databases including L2L (Newman and Weiner, 2005), SignatureDB (Shaffer et al., 2006), CCancer (Dietmann et al., 2010), GeneSigDB (Culhane et al., 2010), GeneSetDB (Araki et al., 2012), and MSigDB (Liberzon et al., 2011). The latter three are widely used for gene set analysis.

MSigDB is the gene set database integrated with GSEA (Subramanian et al., 2005). MSigDB acquires gene sets through manual curation and computational methods (Liberzon et al., 2011). As a meta-database, MSigDB extracts gene sets from several sources including GO (Ashburner et al., 2000), KEGG (Kanehisa et al., 2016), Reactome (Joshi-Tope et al., 2005), and BioCarta (Nishimura, 2001).

GeneSigDB is another database of gene sets extracted from published experimental expression studies of genes, proteins, or miRNAs. GeneSigDB relied on PubMed searches to collect papers relevant to a set of search terms mainly focused on cancer, lung disease, development, immune cells, and stem cells. To develop the database, the authors downloaded the relevant papers and then manually transcribed gene sets from them or their supplementary documents.

GeneSetDB, as another meta-database, is a collection of 26 public databases focused on pathways, phenotypes, drugs, gene regulation, or Gene Ontology. The primary focus of GeneSetDB is human, although it supports mouse and rat using computationally inferred homology (Araki et al., 2012).

### 3.1 Gene set overlap and ORA: A hypothetical example

To show how overlap of gene sets can affect the results of ORA, in this section we present a hypothetical example. Suppose that in a high-throughput experiment, the expression activity of 10000 genes has been measured. After conducting the experiment and performing single gene analysis, 100 genes have been predicted as being differentially expressed. Consider gene sets *A*, *B*, and *C* as illustrated in Figure 2, where gene sets are depicted as circles, and genes belonging to each gene set are depicted as rectangles. In each gene set, genes predicted as being differentially expressed are coloured in red and the rest of the genes are coloured in white. As shown in Figure 2, all genes in *A* have been predicted as being differentially expressed. Table 2 illustrates the contingency table for over-representation of *B*.

**Figure 2:**
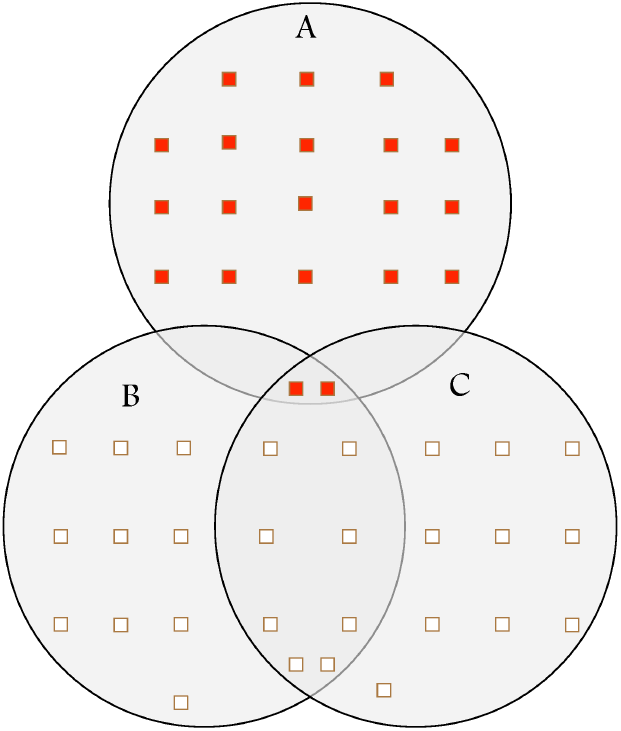
A hypothetical example: gene set overlap leading to lack of specificity of ORA. Each circle represents a gene set. Rectangles coloured in red and white represent differentially expressed and non-differentially expressed genes, respectively. Gene set B (and also C) is predicted as being differentially enriched by ORA solely due to partial overlap with A, a truly differentially enriched gene set.

**Table 2:** The contingency table for the over-representation of *B*. DE stands for differentially expressed and Non-DE stands for non-differentially expressed.

|  | DE Genes | Non-DE Genes | Total |
| --- | --- | --- | --- |
| Genes in B | 2 | 18 | 20 |
| Genes not in B | 98 | 9882 | 9980 |
| Total | 100 | 9900 | 10000 |

According to Fisher’s exact test, *B* is predicted as being differentially enriched with a 95% confidence level (*p-value* = 0.0167). This result is primarily due to the overlap between *A* and *B*. This example suggests that gene set overlap can lead to a lack of specificity in gene set analysis methods.

In this paper, we use GeneSigDB version 4, GeneSetDB for Human (downloaded on February 2, 2018), and MSigDB version 6.0, unless stated otherwise.

### 3.2 Measuring gene set overlap

To study gene set overlap and its effect on the specificity of ORA, we use the Jaccard coefficient to quantify the overlap between two gene sets. We then use this quantitative measure to visualize gene set overlap in MSigDB, GeneSetDB, and GeneSigDB.

For given sets A and B, the Jaccard coefficient is defined as follows:

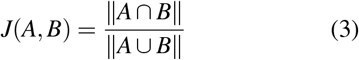

The Jaccard coefficient is a value between 0 and 1, where *J*(*A, B*) = 0 means that there is no overlap between *A* and *B*; *J*(*A, B*) = 1 means that there is a complete overlap between *A* and *B*, i.e. *A* = *B*; and other values (0 < *J*(*A, B*) < 1) represent partial overlaps between *A* and *B*. The Jaccard index can be used to quantify the overlap between two sets; for example, it can be used to measure the overlap between two gene sets from a gene set database or the overlap between a gene set and a set of differentially expressed genes resulting from a gene expression study. Hereafter, we refer to Jaccard index as overlap score.

For a given set of genes *L_i_* and a gene set database 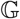, we define the overlap coefficient, or overlap score, of *L_i_* with respect to 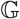 as follows:

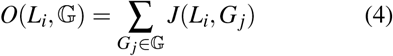

This measure is representative of the cumulative overlap of *L_i_* with all gene sets in the gene set database 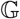. For the sake of brevity, whenever gene set database 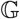 can be inferred from the context, we use the phrase “overlap score of *L_i_*” to refer to 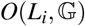. Note that 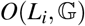, which is the summation of overlap between *L_i_* and each gene set in the gene set database 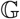, should not be mistaken with overlap between two sets of genes. The latter is calculated using the Jaccard index (Equation 3).

### 3.3 Visualization of gene set overlap

We visualize a gene set database as a graph, where each gene set *G_i_* is represented as a vertex *v_i_*, and there is an edge between two vertices *v_i_* and *v _j_* if *J*(*G_i_, G_j_*) > 0; the value of *J*(*G_i_, G_j_*) is used as the weight for this edge. Since the Jaccard coefficient is symmetric, the graph defined using this measure is an undirected graph. Due to the sheer number of overlapping gene sets, such a graph has a large number of edges. To visualize substantial overlaps between gene sets, we only show overlap scores greater than or equal to 0.5, while retaining all vertices. In other words, in all graph visualizations in this paper, an edge between two vertices *v_i_* and *v _j_* indicates that their corresponding gene sets, i.e. *G_i_* and *G_j_*, share at least half of their genes. The “hairballs” in Figure 3 and also Figures 6 and 7 (in the Appendix) are due to the existence of a large number of edges, i.e. pairs of gene sets with a substantial amount of overlap. These graphs highlight the existence of gene set overlap as a ubiquitous phenomenon in gene set databases. The graph visualization can be generated using Fruchterman Reingold layout (Fruchterman and Reingold, 1991) in Gephi (version 0.9.2) (Bastian et al., 2009).

**Figure 3:**
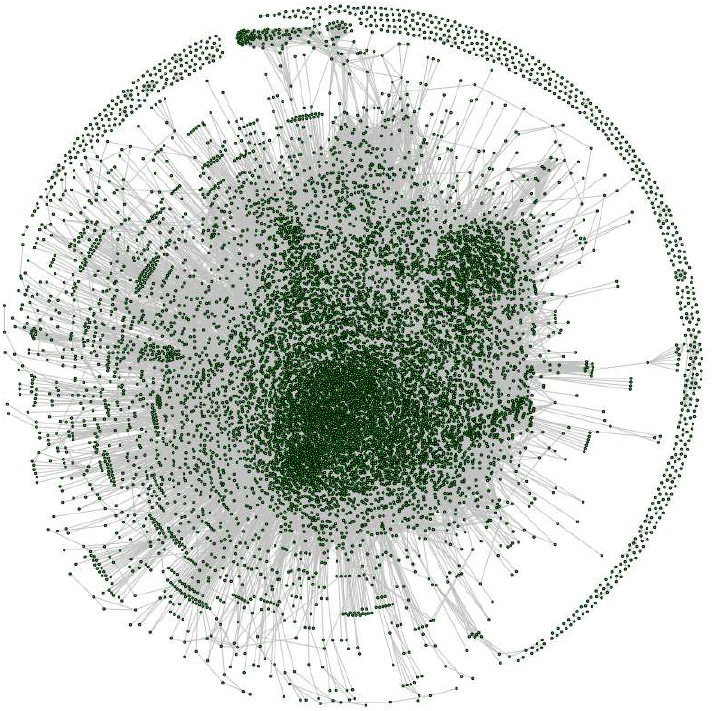
The graph representing the overlap between gene sets in MSigDB. In this graph, each vertex represents a gene set in MSigDB, and each edge represents an overlap with Jaccard coefficient greater than or equal to 0.5 between two gene sets (see Equation 3). The “hairball” is the result of a large number of gene sets with a substantial overlap (≥0.5) with each other.

To further visually inspect the gene set overlap in a given gene set database 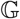, we use a frequency plot. For each gene set *G_i_* in 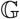, we calculate *f_i_* = ‖{*G_j_* | *J*(*G_i_*, *G_j_*) > 0 (*j* ≠ *i*) and 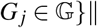. *f_i_* is the number of gene sets *G_j_* (*j* ≠ *i*) in 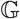 with a non-zero overlap with *G_i_*. After calculating *f_i_* values for all *G_i_* in 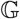, we use a frequency plot to show the distribution of *f_i_* values. Figure 4 and also Figures 8, and 9 (in the Appendix) illustrate the distribution of *f_i_* values for MSigDB, GeneSetDB, and GeneSigDB, respectively. These figures are in agreement with Figure 3, 6, and 7 and show the prevalence of gene set overlap in the aforementioned gene set databases.

**Figure 4:**
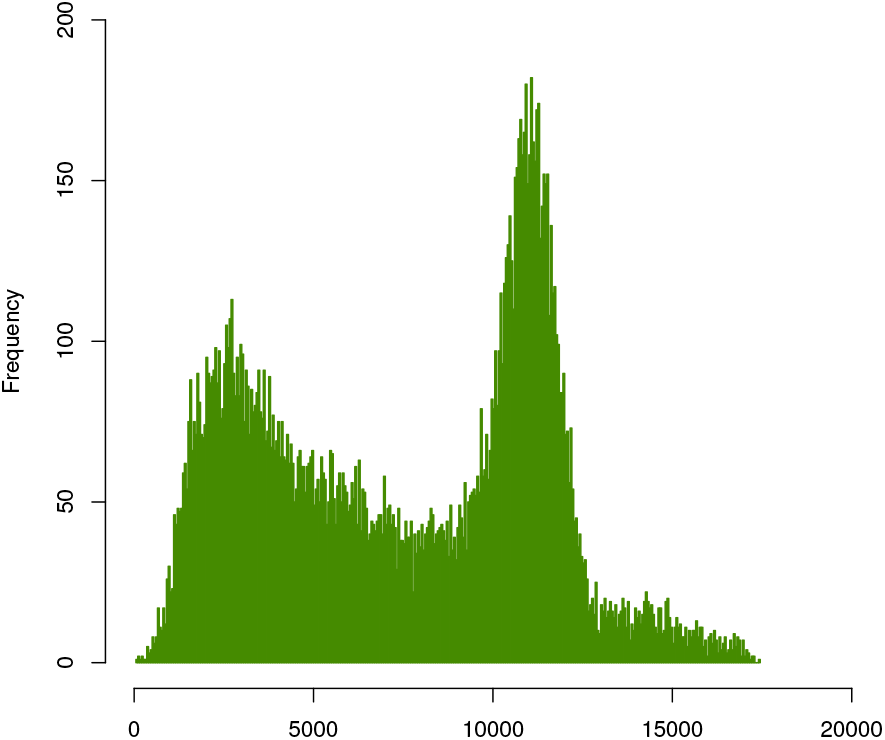
A frequency plot for *f_i_* values in MSigDB illustrates the prevalence of gene set overlap. For each gene set *G_i_* in a gene set database 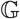 (MSigDB here), *f_i_* is the number of gene sets *G_j_* (*j* ≠ *i*) in 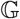 with *J*(*G_i_*, *G_j_*) > 0.

Figure 4 suggests that overlap scores in MSigDB follow a multimodal distribution. This can be attributed to the fact that MSigDB is a meta-database that extracts gene sets from several sources including GO, KEGG, Reactome, and BioCarta. A compelling result revealed by Figure 4 is that majority of gene sets in MSigDB have at least a non-zero overlap with more than 1000 other gene sets in MSigDB (out of a total of 17778 gene sets). Also, there is no gene set in MSigDB without overlap with some other gene set(s). Finally, there are gene sets that overlap with the majority of gene sets in MSigDB. For example, the gene set associated with the “cellular response to organic substance” GO term (GO:0071310) has one non-zero overlap with 17292 gene sets. This gene set is associated with a general GO term and therefore overlaps a large number of gene sets including the gene sets defined using relatively more specific GO terms.

## 4 METHODOLOGY

Evaluation of ORA using a quantitative measure such as specificity requires a gold standard dataset for which the differentially enriched gene sets are *a priori* known. Such a gold standard does not exist. In this section, we propose a methodology for a quantitative evaluation of the effect of gene set overlap on the specificity of ORA in the absence of such a gold standard dataset.

To perform ORA, a single gene analysis method must be conducted to predict the set of differentially expressed genes. This set serves as one of the inputs to ORA. In practice, often noise and biological variability introduce errors—i.e. false positives and false negatives—in the result of single gene analysis. In the context of single gene analysis, false positives are genes that are not differentially expressed but predicted as being so, and false negatives are genes that are differentially expressed but predicted as not being such. False negatives in single gene analysis may reduce the sensitivity of ORA, while false positives may reduce the specificity. To avoid the interference of the single gene analysis errors in the study of gene set overlap and its effect on the specificity of ORA, we assume that differentially expressed genes have been identified correctly; also, this is the same assumption that ORA relies on. Therefore, to perform the quantitative evaluation, a scenario in which all genes in a given gene set have been accurately detected as being differentially expressed is considered.

To deal with the absence of a gold standard dataset, in this paper the following procedure is used to identify the true enrichment status of gene sets. Given a gene set database 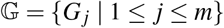 and *L_i_*, a set of differentially expressed genes, and a fixed parameter γ, for each gene set 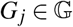 we consider *G_j_* as being truly differentially enriched if at least 100×γ percent of its members are differentially expressed genes, i.e. 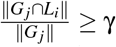. Otherwise, *G_j_* is considered as not being truly differentially enriched. γ serves as a threshold; since there is no consensus about such a threshold value, we repeat the main experiments for a wide range of values for γ, and we show that regardless of the value chosen for γ the results are consistent. In the rest of the paper, the set of truly differentially enriched and truly nondifferentially enriched gene sets are denoted by 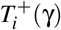 and 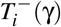, respectively, and are defined as follows:

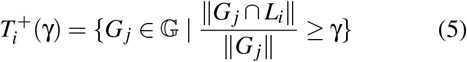

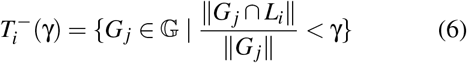

Hereafter, for the sake of brevity, we avoid writing the parameter γ; for example, we refer to 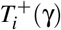 and 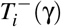 as 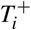 and 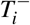 respectively.

Given γ and *L_i_*, Equations 5 and 6 determine the true enrichment status of all gene sets in 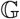. Knowing the true enrichment status of gene sets, we run ORA. The parameters (inputs) for running ORA are: a list of differentially expressed genes *L_i_*, a significance level α, a background set *U*, and a gene set database 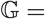 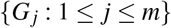.

In this research, the experiments were conducted using *python* version 3.6.2. To implement ORA, the *fisher_exact* method from the *stats* module of *scipy* version 0.19.1 was used. Also, the Benjamini-Hochberg FDR adjustment for multiple comparisons was performed using the *multipletests* method (with method parameter equal to *fdr_bh*) from *statsmodels* version 0.8.0.

For each gene set *G_j_* in 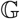, ORA calculates a *p*-value *p _j_*. After calculating *p*_1_*, …, p_m_*—the *p*-values corresponding to the over-representation of gene sets *G*_1_*, …, G_m_* in 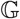 according to Equation 2—the Benjamini-Hochberg FDR adjustment (Drăghici, 2016) for multiple comparisons is applied. All gene sets with an adjusted p-value less than α are predicted as significant, i.e. being differentially enriched. 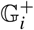 is defined as the set of all such significant gene sets. 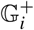 includes both true positives and false positives. 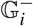 is defined as the set of all nonsignificant gene sets, i.e. 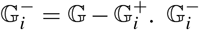 includes both true negatives and false negatives. For the given value of γ, true positives (*TP_i_*), false positives (*FP_i_*), true negatives (*TN_i_*), and false negatives (*FN_i_*) are identified based on Equations 7, 8, 9, and 10.

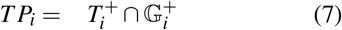

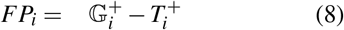

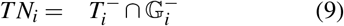

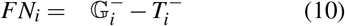

Using these values, specificity (*SPC_i_*) is calculated according to Equation 11.

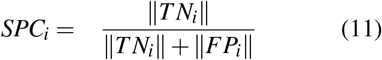

To be able to gain insight that is unbiased toward a single set *L_i_*, this process is repeated many times, each time with a different *L_i_*. We denote the set of all *L_i_* as 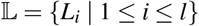.

Algorithm 1 (see the Appendix) illustrates the methodology for conducting the experiment. In each iteration of the algorithm, i.e. the outer loop, a gene set *L_i_* from 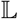 is used, and the process is repeated for all gene sets in 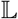. In addition, for each set 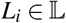, the overlap score of *L_i_* with respect to gene set database 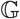, i.e. 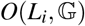, is calculated according to Equation 4. Having overlap score and specificity measure for each 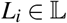, the relationship between overlap and the specificity of ORA can be assessed using statistical methods (see Section 5).

## 5 EXPERIMENTAL RESULTS

To study the effect of gene set overlap on the specificity of ORA using Algorithm 1, MSigDB—one of the most widely used gene set databases devoted to gene set analysis—was used as the gene set database 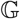. Since ORA requires a list (set) of differentially expressed genes as input, Algorithm 1 requires a collection of such lists (denoted as 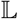 in the algorithm). ImmuneSigDB (Godec et al., 2016) version 6.0 was used to provide such a collection. ImmuneSigDB contains lists of differentially expressed genes, each created by identifying differentially expressed genes in a dataset extracted from Gene Expression Omnibus (GEO) (Edgar et al., 2002). Therefore, each list in ImmuneSigDB represents a set of differentially expressed genes derived from a high-throughput study.

To investigate the association between gene set overlap and the specificity of ORA results, first the overlap score 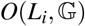 was calculated for each list *L_i_* in ImmuneSigDB. In this experiment, a significance level α = 0.05 and γ values equal to 0.1, 0.2, 0.3, 0.4, 0.5, 0.6, 0.7, 0.8, 0.9, and 0.99 were used. For each value of γ, Algorithm 1 was run to calculate *SPC_i_* corresponding to each 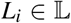. Figure 5 illustrates the relationship between gene set overlap and the number of false positives for γ = 0.5. As overlap score increases, we observe an increase in the number of false positives and therefore a decline in the specificity. We observed the same pattern for all the aforementioned values of γ.

**Figure 5:**
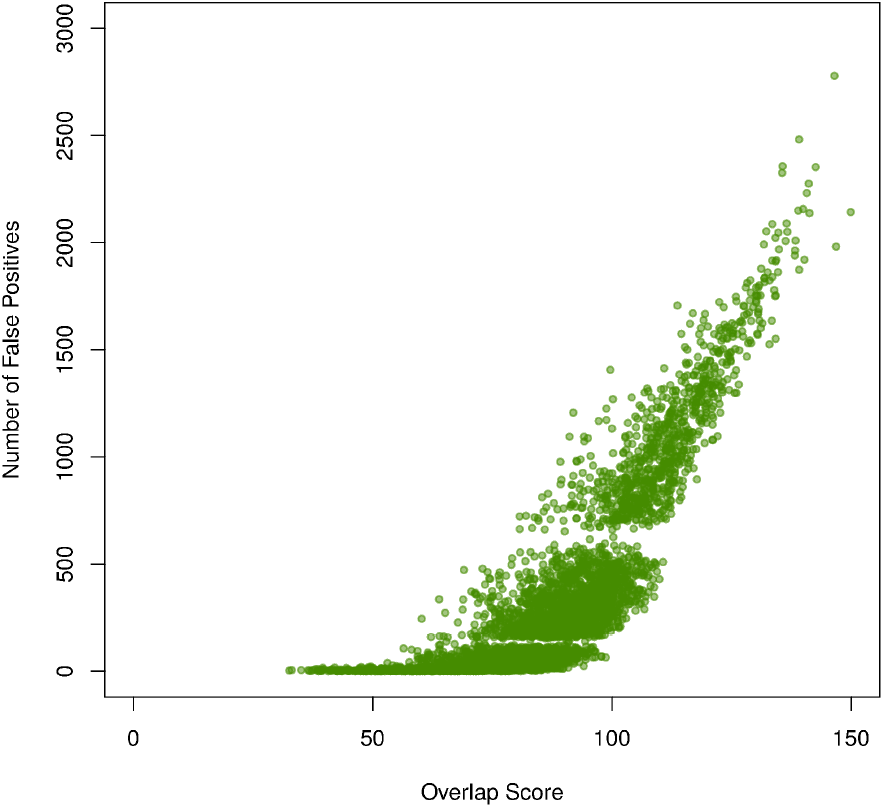
Number of false positives increases as overlap score increases (γ = 0.5). A similar pattern was observed for other values of γ.

To study the relationship between gene set overlap and the specificity of ORA, we used a statistical test of correlation. Choosing a proper test of correlation requires assessment of the normality assumption. To test the null hypothesis that specificity values are normally distributed, we used the Shapiro-Wilk test (Shapiro and Wilk, 1965). Table 4 shows the test results for the aforementioned values of γ. Considering those results, as confirmed by the histogram in Figure 10 (see Appendix), we concluded that specificity values are not normally distributed. Therefore, a Spearman’s rank correlation coefficient test, a non-parametric test, was conducted for each value of γ to test the null hypothesis that there is no correlation between specificity and overlap scores. Table 3 shows the result of this test for various values of γ. Considering these results, we concluded that there is a strong negative correlation between gene set overlap and specificity of ORA.

**Table 3:**
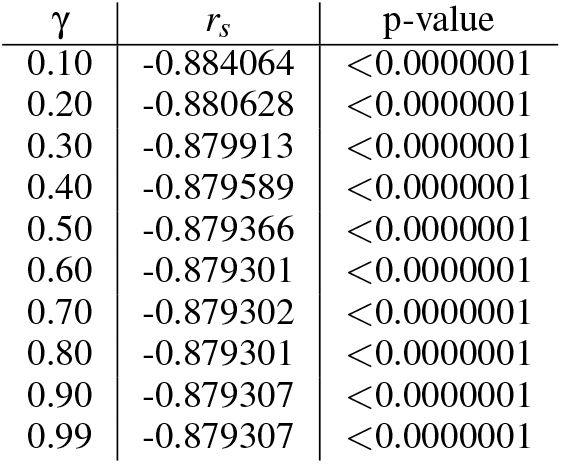
The result of Spearman rank correlation tests for different values of γ. All p-values are less than 0.0000001.

| $\gamma$ | $r_s$ | p-value |
| --- | --- | --- |
| 0.10 | -0.884064 | <0.0000001 |
| 0.20 | -0.880628 | <0.0000001 |
| 0.30 | -0.879913 | <0.0000001 |
| 0.40 | -0.879589 | <0.0000001 |
| 0.50 | -0.879366 | <0.0000001 |
| 0.60 | -0.879301 | <0.0000001 |
| 0.70 | -0.879302 | <0.0000001 |
| 0.80 | -0.879301 | <0.0000001 |
| 0.90 | -0.879307 | <0.0000001 |
| 0.99 | -0.879307 | <0.0000001 |

## 6 DISCUSSION

In this research, we proposed a systematic approach for evaluating the specificity of over-representation analysis. Using the proposed method, we demonstrated that there is a significant negative correlation between the specificity of ORA and gene set overlap. In other words, gene set overlap increases the number of false positives, i.e. gene sets incorrectly predicted as being differentially enriched. The increase in the number of false positives makes interpreting the results of ORA difficult and prone to investigator biases toward a hypothesis of interest. It also hinders reproducibility of gene set analysis results.

We also showed that gene set overlap is a ubiquitous phenomenon across gene set databases. The existence of multifunctional genes is one contributor to this phenomenon. Multifunctional genes are genes associated with several molecular functions or biological processes; therefore, they appear in several gene sets, contributing to gene set overlap. Multi-functional genes are commonplace; for example, Pritykin et al. (Pritykin et al., 2015) identified 2517 multifunctional genes in the human genome. As a consequence, gene set overlap is an integral characteristic of gene set databases. Another factor contributing to the prevalence of gene set overlap in databases that define some (or all) of their gene sets based on GO is the child-parent relationship between GO terms. GO terms are organized as a directed acyclic graph; each node represents a GO term; and each edge between two nodes represents a parent-child relationship between terms, with the child term being more specific than its parent term(s). Therefore, gene sets derived from GO terms that are involved in such child-parent relationships share common genes; this, in turn, contributes to the existence of gene set overlap.

Being an integral part of gene set databases, gene set overlap should be considered in the design and evaluation of gene set analysis methods. However, many gene set analysis studies have used simulated collections of non-overlapping gene sets for method evaluation and comparison (Ackermann and Strimmer, 2009), (Efron and Tibshirani, 2007), (Nam and Kim, 2008). Therefore, gene set overlap and its effect on the outcome of gene set analysis methods have been overlooked. We suggest using datasets that account for overlap as a requirement in the evaluation of gene set analysis methods.

Although many gene set analysis methods and tools have been developed, there are very few methods that consider gene set overlap. For example, PADOG is an attempt for addressing gene set overlap that leads to a small number of false positives (has high specificity), but its sensitivity has been reported to be lower than that of other gene set analysis methods (see Table S2 from the work by Tarca et al., 2013). SetRank is another gene set analysis method designed with gene set overlap in mind to increase specificity (Simillion et al., 2017). The authors of SetRank claimed that due to a lower number of false positives, the significant results reported by this method are more reliable than other methods. Therefore, it may be a viable solution for the lack of specificity of gene set analysis methods. A rigorous evaluation of the specificity and sensitivity of this method is suggested as future research.

The existence of gene set databases that accurately represent biological processes and functions is essential to the success of gene set analysis. Increasing the size of gene set databases by depositing more gene sets has been the common trend in developing gene set databases. The increase in the number of gene sets has introduced more gene set overlap, which in turn leads to a higher false positive rate. There is a need to focus on quality rather than sheer quantity in developing gene set databases. We suggest further research on the quality control of gene set databases.

Another suggestion for improving the specificity of current methods is to exclude irrelevant or uninformative gene sets before conducting gene set analysis. Considering the size of gene set databases, filtering these gene sets is laborious and, if done manually, prone to investigator bias toward gene sets considered “relevant”. Developing a computational approach for filtering irrelevant or uninformative gene sets would be worthwhile.

In the proposed method for evaluating ORA, we considered scenarios with only one differentially enriched gene set. In practice, a specific phenotype may be the result of altering several biological processes or functions, i.e. multiple gene sets. We expect that the differential enrichment of several gene sets intensifies the extent to which gene set overlap reduces specificity. In other words, we expect to see a larger number of false positives compared to the situation considered in this work. The proposed method is capable of handling scenarios with several differentially enriched gene sets. Also, Algorithm 1 can be used seamlessly with sensitivity or accuracy instead of specificity.

Since the input to ORA is a list of differentially expressed genes, we utilized ImmuneSigDB (Godec et al., 2016) for evaluating ORA. However, some gene set analysis methods require an expression matrix that represents expression level of genes under study across control and case samples. The proposed methodology is capable of evaluating such gene set analysis methods. To do so, the only requirement is developing expression profiles with the differentially enriched gene set(s) encoded in expression values. Therefore, our methodology can be used as a systematic approach to study specificity, sensitivity, and accuracy of other gene set analysis methods. For example, we suggest the study of the relationship between gene set overlap and the specificity of GSEA (Subramanian et al., 2005), which is another well-established gene set analysis method, as future work.

In the absence of gene set overlap, gene set analysis is a trivial problem, as many methods have achieved high specificity when being evaluated (by their authors) using simulated gene set databases with non-overlapping gene sets. If gene set overlap was considered in the evaluation of these methods, the lack of specificity of many gene set analysis methods would be obvious. For example, assume a gene set analysis method that uses average expression value of genes within a gene set (in control versus case samples) to predict the enrichment status of a gene set. Also assume that there is a single differentially expressed gene that appears in 100 gene sets. Such a method would report all 100 gene sets as being differentially enriched, while most of these gene sets might be biologically irrelevant. Therefore, we strongly recommend considering gene set overlap in any attempt for evaluating gene set analysis using simulated data.

## 7 CONCLUSION

In this paper, we proposed a systematic approach to study the effect of gene set overlap on the result of ORA (over-representation analysis). Using the proposed method and statistical analysis, we showed that there is a significant negative correlation between gene set overlap and specificity of ORA. We quantified gene set overlap and showed that it is a ubiquitous phenomenon across gene set databases. The proposed approach for the study of the relationship between gene set overlap and specificity of ORA can easily be used to investigate the effect of gene set overlap on different gene set analysis methods using quantitative measures such as specificity, sensitivity, and accuracy.

Considering the effect of gene set overlap on the results of ORA, it is essential to develop and use methods that address gene set overlap and achieve higher specificity without sacrificing sensitivity in the prediction of differentially enriched gene sets. Due to the lack of gold standard datasets, where the differentially enriched gene sets are known *a priori*, simulated datasets have been widely used for evaluation of gene set analysis methods. The databases used in these studies are often a collection of non-overlapping gene sets of the same size. This setting is substantially different from a real gene set database where gene set overlap is common. By completely ignoring gene set overlap, some methods achieve high specificity on simulated data but behave inadequately when working in real settings. We strongly recommend that the use of non-overlapping datasets be avoided for evaluation of gene set analysis methods.

## APPENDIX

**Algorithm 1.**
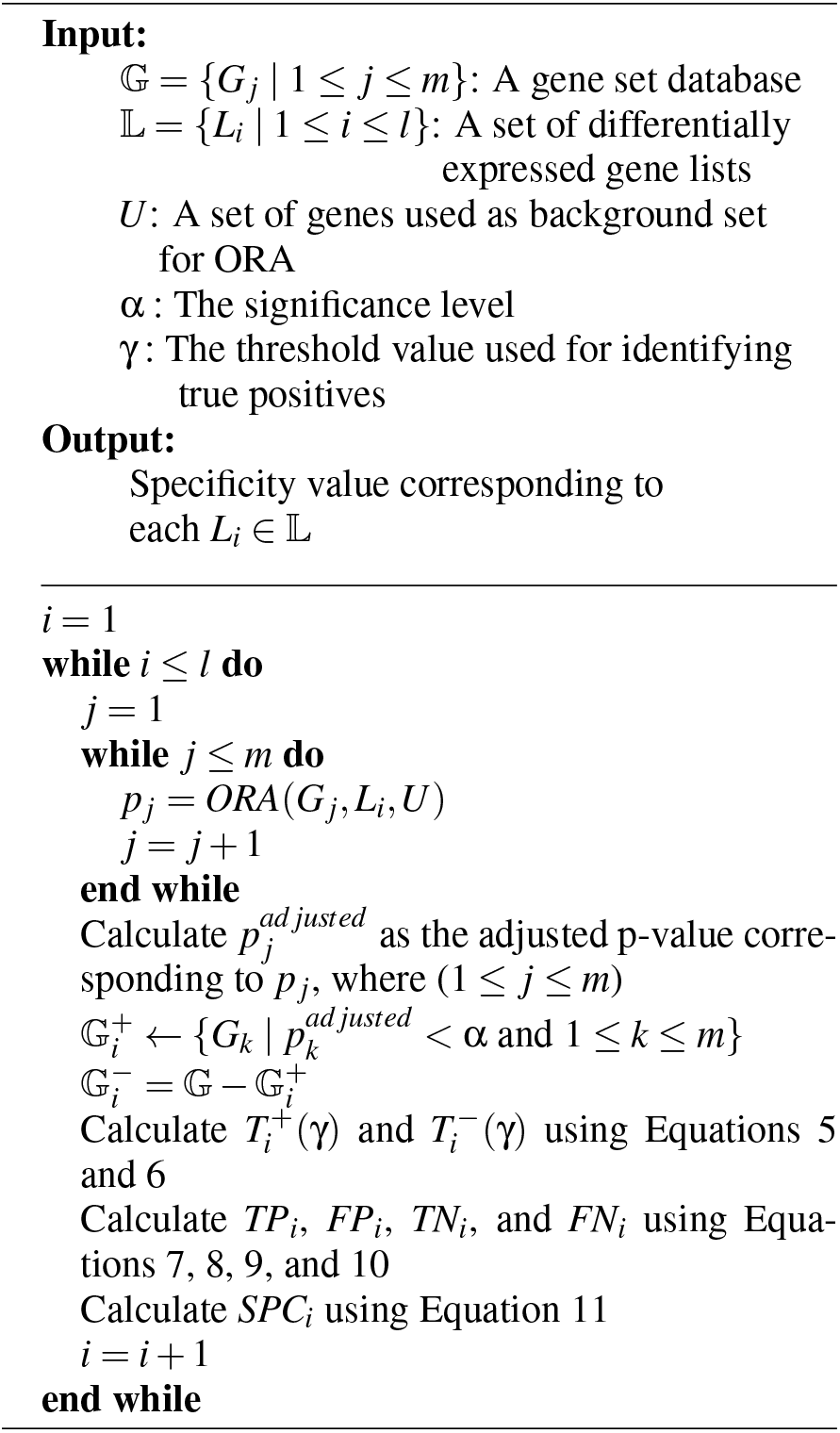
Calculation of specificity of ORA

**Table 4:**
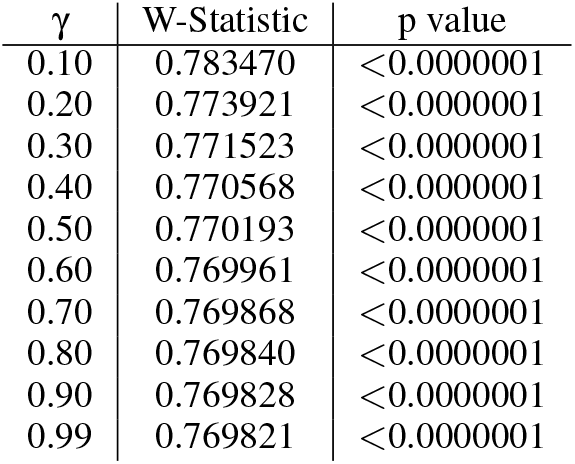
The result of Shapiro-Wilk tests for different values of γ. All p-values are less than 0.0000001.

**Figure 6:**
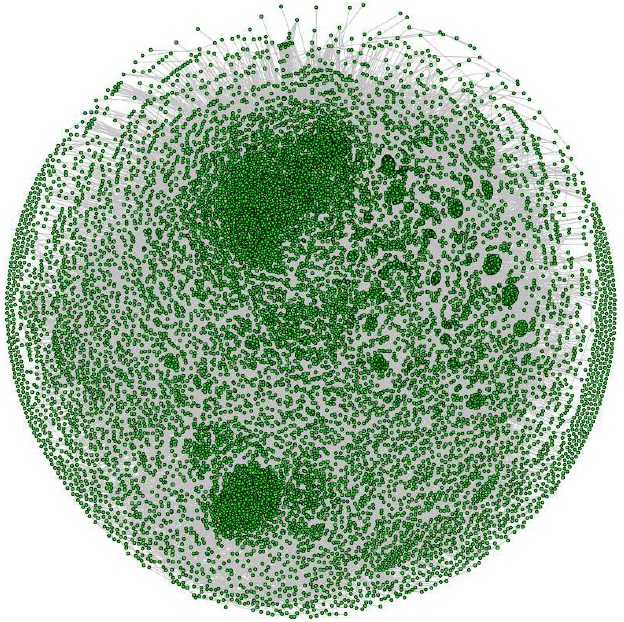
The graph representing the overlap between gene sets in GeneSetDB. In this graph, each vertex represents a gene set in GeneSetDB, and each edge represents an overlap with Jaccard coefficient greater than or equal to 0.5 between two gene sets. The “hairball” is the result of a large number of gene sets with a substantial overlap (≥0.5) with each other.

**Figure 7:**
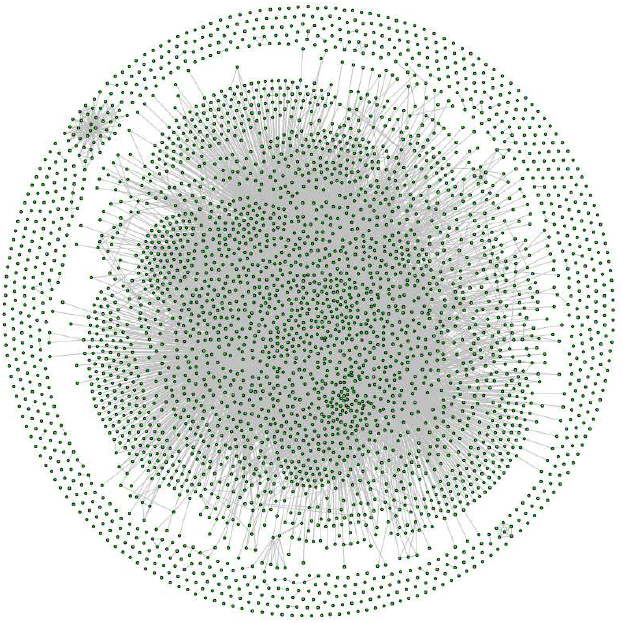
The graph representing the overlap between gene sets in GeneSigDB. In this graph, each vertex represents a gene set in GeneSigDB, and each edge represents an overlap with Jaccard coefficient greater than or equal to 0.5 between two gene sets. The “hairball” is the result of a large number of gene sets with a substantial overlap (≥0.5) with each other.

**Figure 8:**
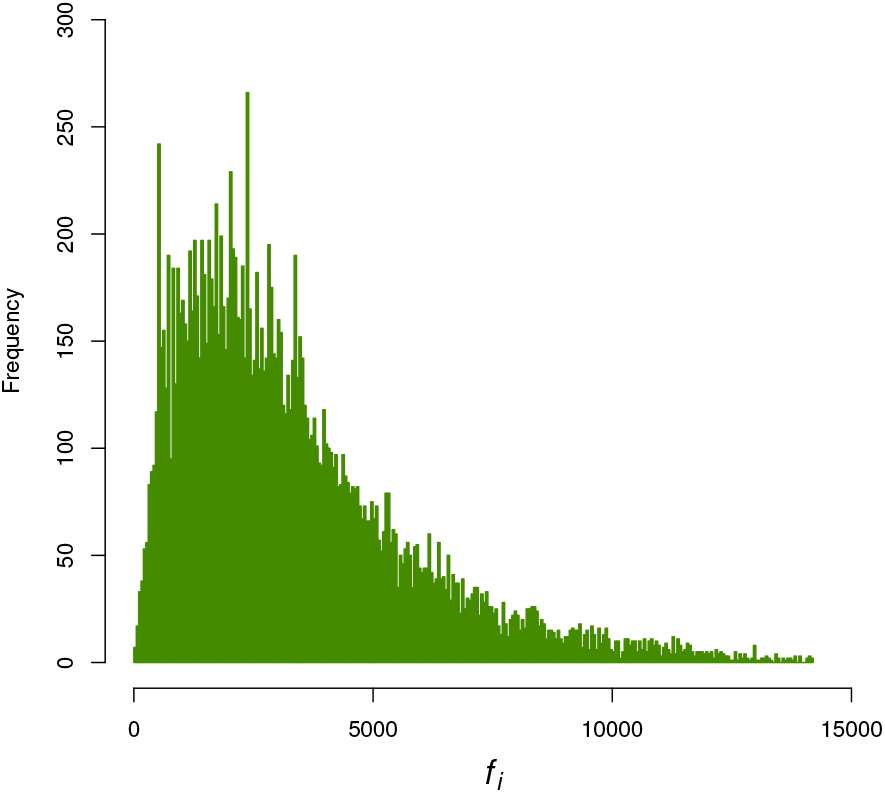
A frequency plot for *f_i_* values in GeneSetDB illustrates the prevalence of gene set overlap. For each gene set *G_i_* in a gene set database 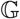 (GeneSetDB here), *f_i_* is the number of gene sets *G_j_* (*j* ≠ *i*) in 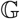 with a non-zero overlap with *G_i_*.

**Figure 9:**
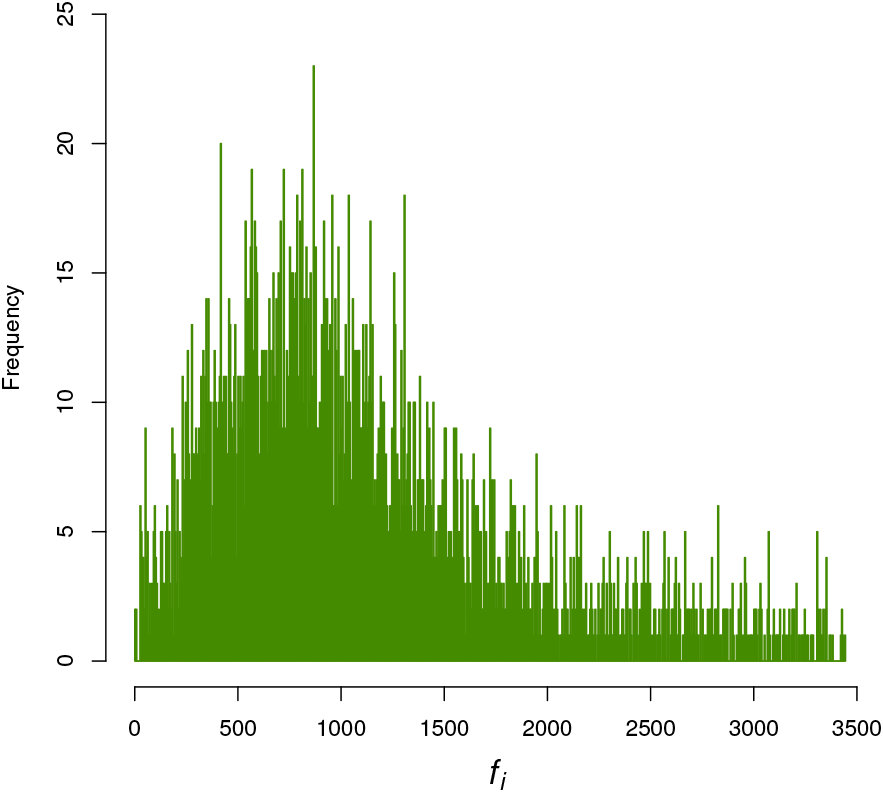
A frequency plot for *f_i_* values in GeneSigDB illustrates the prevalence of gene set overlap. For each gene set *G_i_* in a gene set database 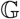 (GeneSigDB here), *f_i_* is the number of gene sets *G_j_* (*j* ≠ *i*) in 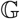 with a non-zero overlap with *G_i_*.

**Figure 10:**
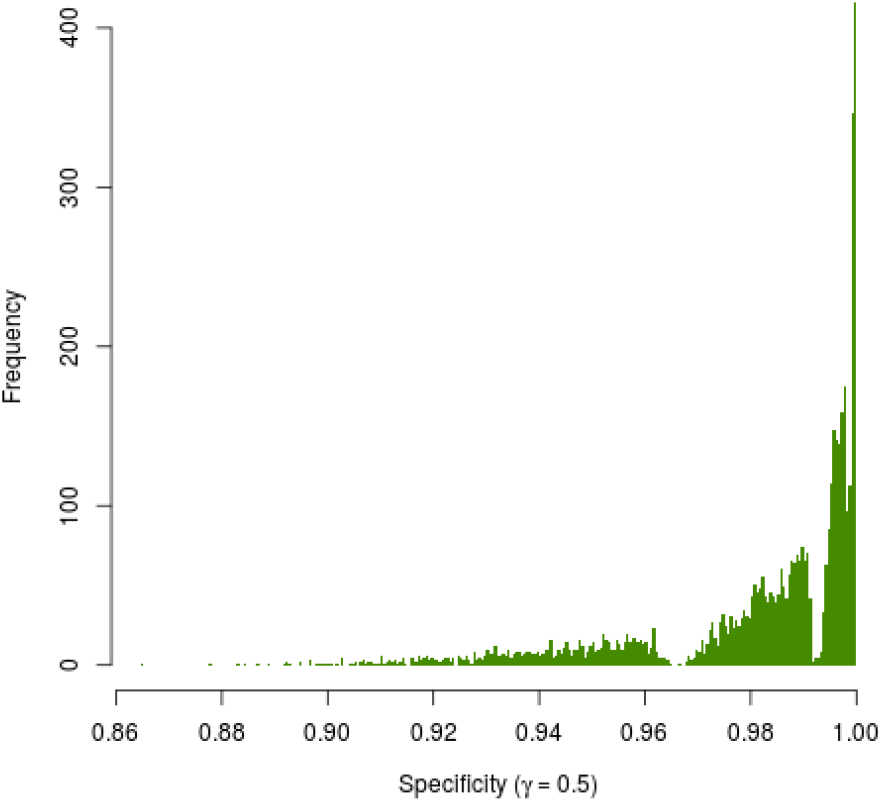
The histogram of the specificity values (γ = 0.5). Obvious deviation of the histogram from a bell-shaped curve suggests that the specificity values are not normally distributed. A similar pattern was observed for other values of γ.

## Notes

### Competing Interest Statement

The authors have declared no competing interest.

### Summary of Updates

The paper has been revised to reflect the changes applied before submission to the 12th International Joint Conference on Biomedical Engineering Systems and Technologies.

