## Appendix for "Gene Set Overlap: An Impediment to Achieving High Specificity in Over-representation Analysis"

**Algorithm 1** Calculation of specificity of ORA

**Input:**
$\mathbb{G} = \{G_j \mid 1 \le j \le m\}$: A gene set database
$\mathbb{L} = \{L_i \mid 1 \le i \le l\}$: A set of differentially expressed gene lists
$U$: A set of genes used as background set for ORA
$\alpha$ : The significance level
$\gamma$ : The threshold value used for identifying true positives

**Output:**
Specificity value corresponding to each $L_i \in \mathbb{L}$

```
i = 1
while i ≤ l do
    j = 1
    while j ≤ m do
        p_j = ORA(G_j, L_i, U)
        j = j + 1
    end while
    Calculate p_j^adjusted as the adjusted p-value corresponding to p_j, where (1 ≤ j ≤ m)
    G_i^+ ← {G_k | p_k^adjusted < α and 1 ≤ k ≤ m}
    G_i^- = G − G_i^+
    Calculate T_i^+(γ) and T_i^-(γ) using Equations 5 and 6
    Calculate TP_i, FP_i, TN_i, and FN_i using Equations 7, 8, 9, and 10
    Calculate SPC_i using Equation 11
    i = i + 1
end while
```

Table 4: The result of Shapiro-Wilk tests for different values of $\gamma$. All p-values are less than 0.0000001.

| $\gamma$ | W-Statistic | p value |
|---|---|---|
| 0.10 | 0.783470 | <0.0000001 |
| 0.20 | 0.773921 | <0.0000001 |
| 0.30 | 0.771523 | <0.0000001 |
| 0.40 | 0.770568 | <0.0000001 |
| 0.50 | 0.770193 | <0.0000001 |
| 0.60 | 0.769961 | <0.0000001 |
| 0.70 | 0.769868 | <0.0000001 |
| 0.80 | 0.769840 | <0.0000001 |
| 0.90 | 0.769828 | <0.0000001 |
| 0.99 | 0.769821 | <0.0000001 |

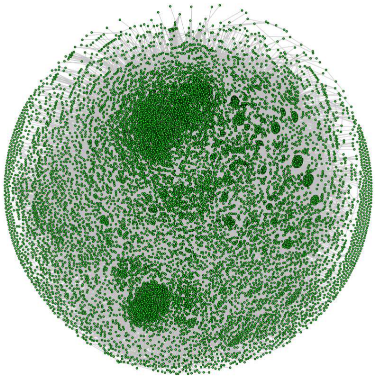

Figure 6: The graph representing the overlap between gene sets in GeneSetDB. In this graph, each vertex represents a gene set in GeneSetDB, and each edge represents an overlap with Jaccard coefficient greater than or equal to 0.5 between two gene sets. The "hairball" is the result of a large number of gene sets with a substantial overlap ($\geq 0.5$) with each other.

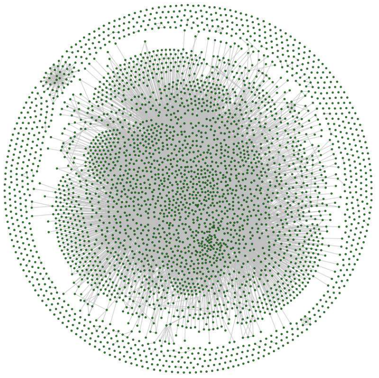

Figure 7: The graph representing the overlap between gene sets in GeneSigDB. In this graph, each vertex represents a gene set in GeneSigDB, and each edge represents an overlap with Jaccard coefficient greater than or equal to 0.5 between two gene sets. The "hairball" is the result of a large number of gene sets with a substantial overlap ($\geq 0.5$) with each other.

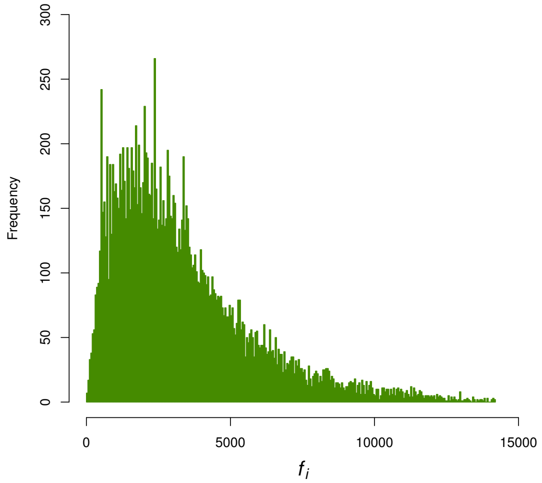


Figure 8: A frequency plot for $f_i$ values in GeneSetDB illustrates the prevalence of gene set overlap. For each gene set $G_i$ in a gene set database $\mathbb{G}$ (GeneSetDB here), $f_i$ is the number of gene sets $G_j$ ($j \neq i$) in $\mathbb{G}$ with a non-zero overlap with $G_i$.

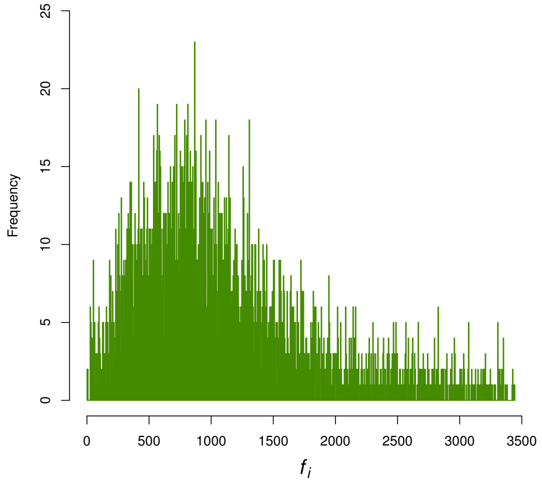


Figure 9: A frequency plot for $f_i$ values in GeneSigDB illustrates the prevalence of gene set overlap. For each gene set $G_i$ in a gene set database $\mathbb{G}$ (GeneSigDB here), $f_i$ is the number of gene sets $G_j$ ($j \neq i$) in $\mathbb{G}$ with a non-zero overlap with $G_i$.

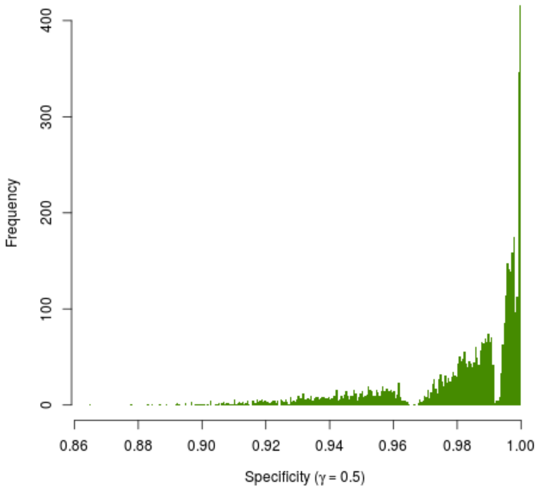


Figure 10: The histogram of the specificity values ($\gamma = 0.5$). Obvious deviation of the histogram from a bell-shaped curve suggests that the specificity values are not normally distributed. A similar pattern was observed for other values of $\gamma$.
